# Deletion of the *Saccharomyces cerevisiae RACK1* homolog, *ASC1*, enhances autophagy which mitigates TDP-43 toxicity

**DOI:** 10.1101/2025.09.05.674500

**Authors:** Sei-Kyoung Park, Sangeun Park, Susan W. Liebman

**Affiliations:** Department of Pharmacology, University of Nevada, Reno, 1664 N Virginia, Mail stop 318, Reno, NV 89557

**Keywords:** TDP-43 (TAR DNA-binding protein 43), FUS (fused in sarcoma), autophagy, TOROID, ASC1/RACK1

## Abstract

Cytoplasmic aggregation of nuclear proteins such as TDP-43 (TAR DNA-binding protein 43) and FUS (fused in sarcoma) is associated with several neurodegenerative diseases. Studies in higher cells suggest that these aggregates of TDP-43 and FUS sequester polysomes by binding RACK1 (receptor for activated C kinase 1), a ribosomal protein, thereby inhibiting global translation and contributing to toxicity. But RACK1 is also a scaffold protein with many other roles including a role in autophagy. Using yeast we find that deletion of the *RACK1* ortholog, *ASC1*, reduces TDP-43 toxicity, but not FUS toxicity. TDP-43 foci remain liquid like in the presence *asc1Δ* but they become smaller. This is consistent with the findings in cell culture. However, using double label tags we establish that ASC1 does not co-localize with TDP-43 foci, arguing against the sequestration hypothesis. Instead, *ASC1* appears to influence toxicity through autophagy. We previously showed that expression of TDP-43 inhibits autophagy and TOROID (TORC1 Organized in Inhibited Domains) formation and that modifiers that rescue yeast from TDP-43 toxicity reverse these inhibitions. Here we show that FUS does not inhibit autophagy. This autophagy enhanced by *asc1Δ* is non-canonical, marked by reduced TOROID formation, and effectively counteracts the autophagy inhibition caused by TDP-43. Our findings suggest that ASC1 influences TDP-43 toxicity through autophagy regulation rather than polysome sequestration, highlighting autophagy as a key therapeutic target.

**Summary:** TDP-43 and FUS aggregates are linked to neurodegenerative diseases. RACK1, a ribosomal protein, was previously thought to contribute to toxicity by co-localizing with these aggregates and sequestering polysomes. In yeast, deletion of *ASC1*—the *RACK1* homolog—reduces TDP-43 toxicity but not FUS toxicity. TDP-43 foci remain liquid-like, but ASC1 does not co-localize with them, challenging the sequestration hypothesis. Instead, *asc1Δ* enhances autophagy, rescuing cells from the autophagy inhibition caused by TDP-43. Unlike TDP-43, FUS does not inhibit autophagy. These findings highlight autophagy, rather than polysome sequestration, as the key mechanism of TDP-43 toxicity and its mitigation via ASC1/RACK1 reduction.

## Introduction

Cytoplasmic aggregation of nuclear proteins such as TDP-43 (TAR DNA-binding protein 43) and FUS (fused in sarcoma) is associated with several neurodegenerative human diseases including amyotrophic lateral sclerosis (ALS), frontotemporal degeneration (FTD) and limbic-predominant age-related TDP-43 encephalopathy (LATE) (Dormann and Haass 2011). When expressed in *Saccharomyces cerevisiae*, TDP-43 (Johnson *et al*. 2008; Armakola *et al*. 2011) and FUS form cytoplasmic foci and cause toxicity, modeling aspects of human disease (Fushimi *et al*. 2011; Ju *et al*. 2011).

Since TDP-43 and FUS do not have yeast homologs, the yeast model can only be used to study their toxic gain-of-function. Although loss of TDP-43 function also contributes to toxicity in humans (Ma *et al*. 2022; Zeng *et al*. 2024), there is evidence that the gain-of-function TDP-43 toxicity seen in yeast is relevant to human disease. Indeed, mutations in *PBP1* that reduced TDP-43 toxicity in yeast revealed a new genetic risk factor for ALS in humans, intermediate-length polyQ expansions in the PBP1 human homolog, ataxin-2, now screened for in the clinic (Gispert *et al*. 2012; Gitler 2012; Becker *et al*. 2017).

Expression of TDP-43 in yeast inhibits the formation of TORC1 organized in inhibited domains (TOROIDs), thereby activating TOR1, and suppressing autophagy (Park *et al*. 2022). This TDP-43 inhibition of TOROIDs and autophagy is mitigated by the modifiers, *pbp1Δ*, *tip41Δ,* or *aah1Δ apt1Δ,* which also reduce TDP-43 toxicity (Park *et al*. 2024; Park *et al*. 2025).

While the toxic form of TDP-43 in humans is still debated (Capitini *et al*. 2021; Cascella *et al*. 2023; Shenoy *et al*. 2023), TDP-43 foci formed in yeast do not stain with thioflavin T and so are not amyloid-like. In contrast, FUS foci do stain with thioflavin T. Also, while TDP-43 foci are liquid-like droplets that can be dissolved by hexanediol and exhibit dynamic properties, FUS aggregates are resistant to hexanediol. TDP-43 remains liquid like whether it is expressed in strains with or without the *pbp1Δ* or *tip41Δ* modifiers (Park *et al*. 2024). In contrast, *aah1Δ apt1Δ,* which likewise reduces TDP-43 toxicity, converts liquid-like TDP-43 into amyloid-like foci identifying liquid-like foci as the toxic form of TDP-43 in yeast (Park *et al*. 2025).

Receptor for activated C kinase 1 (RACK1) has recently been identified as a modifier of TDP-43 toxicity in higher cells. Knockdowns of RACK1 were reported to decrease cytoplasmic aggregation of TDP-43 and FUS and reduce TDP-43-associated neurodegeneration in transgenic Drosophila melanogaster expressing hTDP-43 (Zhao *et al*. 2023). RACK1/ASC1 sometimes behaves as a 40S ribosomal protein, is involved in signaling, translation, cancer, inflammation, autophagy, etc., and interacts with TDP-43 in a 2-hybrid assay (Adams *et al*. 2011; Dengjel *et al*. 2012; Wolf and Grayhack 2015) (http://www.interactome-atlas.org/).

Importantly, RACK1 was reported to co-aggregate with TDP-43 cytoplasmic foci in a cell line and motor neurons. This led to the novel hypothesis that the pathogenic effect of reducing translation that is associated with TDP-43 aggregates results from sequestration of polysomes due to TDP-43 binding to the small ribosomal protein RACK1 (Russo *et al*. 2017). Similar results have now also been reported for FUS (Zhao *et al*. 2023).

Since the yeast protein ASC1 is orthologous to RACK1 (Gerbasi *et al*. 2004) we examined the effect of ASC1 on TDP-43 and FUS toxicity in yeast. We found that *asc1Δ* reduces TDP-43 toxicity but not FUS toxicity. Also, we did not observe sequestration of ASC1-GFP in foci in the presence of TDP-43. Rather, our results suggest that the increase in autophagy caused by *asc1Δ* (Gross *et al*. 2019) rescues cells from TDP-43-mediated autophagy inhibition (Park *et al*. 2022; Park *et al*. 2024; Park *et al*. 2025).

## Methods

### Strains and plasmids

All experiments were done in *S. cerevisiae* BY4741 ([*PIN*^+^] *MAT***a** *his3Δ1 leu2Δ0 met15Δ0 ura3Δ0*). BY4741 *ASC1* deletion strain (*asc*1Δ) was obtained from Open Biosystems (Huntsville, AL, USA, Cat # YSC1053). To generate plasmid p*PASC1*-*ASC1-mCFP* (p2749) 1) backbone plasmid p*AG413-GAL*-*ccdB-Cerulean* (Addgene plasmid #14381 *CEN, HIS3*) was digested with *SacI* and *EcoRV* to remove the *GAL* promoter and the digested vector was purified via gel extraction; 2) the *ASC1* gene under its own promoter was PCR-amplified from plasmid pEAW076 (p2733) (Wolf and Grayhack 2015), kindly received from Dr. Elizabeth J Grayhack, using Vent DNA polymerase (New England BioLabs Inc.). Forward (5′-CCGGAGCCTcatatgtgatccctggaa-3′) primer included an additional nucleotide to maintain the correct reading frame and reverse (5′-AGGCATATCaaggtattcagagacttacaat-3′) primer lacked a stop codon to allow in-frame fusion with the downstream cerulean coding sequence. The primers were designed to introduce a *SacI* site at the 5′ end and an *EcoRV* site at the 3′ end. The PCR products were digested with *SacI* and *EcoRV*, gel-purified, and ligated into the prepared vector backbone. The resulting constructs were verified by restriction digestion. Plasmid *pGAL-ASC1-*Cerulien (p2738), a *2µ* vector expressing ASC1 from a GAL promoter was constructed using Gateway LR recombination. The destination vector pAG425-GAL-ccdB-Cerulean (p2731) (Addgene plasmid #14393; 2µ, *LEU2*) was used to transfer *ASC1* from its entry clone p2761 (pDONR221-ASC1, KanR), which lacked a stop codon. *ASC1* was PCR-amplified from the *S. cerevisiae* genome using primers: Forward 5’-GGGGACAAGTTTGTACAAAAAAGCAGGCTTAGAAGGAGATAGAACCATGGCATCTAACGAAGTTTTAG -3′) and reverse (5′-GGGGACCACTTTGTACAAGAAAGCTGGGTCGT TAGCAGTCATAACTTGCCAAACTCTAATGACGTTG -3′). The TDP-43 expression plasmids p*GAL*-*TDP-43* (*CEN, LEU2*) (p2368, pAG415-*GAL-TDP-43*) and p*GAL*-*TDP-43* (*CEN, URA3*) (p2665, pAG416-*GAL-TDP-43*) were constructed via Gateway LR recombination using the entry clone p2273 (pDONR221-TDP-43, containing a stop codon) and the destination vectors pAG415-*GAL-ccdB* (Addgene plasmid #14145; *CEN, LEU2*) or pAG416-*GAL-ccdB* (Addgene plasmid #14147; *CEN, URA3*), respectively. These plasmids were previously described (Park *et al*. 2017). Similarly, the pGAL-FUS expression plasmid (p2764, pAG413-*GAL-FUS*; *CEN, HIS3*) was generated by Gateway LR recombination between the destination vector pAG413-*GAL-ccdB* (Addgene plasmid #14141; *CEN, HIS3*) and the entry clone p2763 (pDONR221-FUS, containing a stop codon). pRS416-GAL1-TDP-43-YFP (*CEN, URA3*, Addgene plasmid #27447) and pGAL*-FUS-EYFP* (*CEN*, *URA3* Addgene plasmid #29593) was purchased at Addgene. The plasmid pGFP-ATG8 (p2571, *pCUP*1, *URA3, CEN*, Addgene plasmid #49423, deposited by Daniel Klionsky) was used to measure autophagy.

An N-terminal GFP-tagged TOR1 *asc1Δ strain* was generated by integrating SpeI-digested pSK108 plasmid into the *asc1Δ* background. The pSK108 construct was originally generated by Dr. Robbie Joséph Loewith and kindly provided as a gift (Prouteau *et al*. 2017). Yeast transformations were performed using the lithium acetate method as described (Gietz and Woods 2002). Transformants were selected and cultured on plasmid-selective glucose medium.

### Measuring Toxicity

Transformants were cultured in plasmid-selective synthetic media containing 2% dextrose and all amino acids except those required for plasmid selection. Cultures were normalized to an OD₆₀₀ of 4, and 10-fold serial dilutions were prepared in water. Diluted cells were spotted onto plasmid-selective media containing either 2% glucose or 2% galactose supplemented with 1% raffinose. Plates were incubated at 30 °C. Plates with glucose media were and scanned after 3 days and those with galactose media after 6–7 days.

To quantify viable cells while avoiding suppressor mutants that reverse TDP-43 toxicity, fresh transformants were patched and maintained on dextrose-selective media, then replica-plated onto plasmid-selective galactose media. For microscopic cell imaging, TDP-43-expressing cells were resuspended in 10x volume of TE buffer. A 1.5 μL aliquot of the cell suspension was mixed with 1.5 μL of 0.4% trypan blue on a microscope slide. Cells were imaged using a Nikon Eclipse E600 fluorescence microscope (Tokyo, Japan) equipped with 100×/1.23 NA or 60×/1.4 NA oil immersion objectives. Fields were photographed, and cells expressing TDP-43-YFP were scored for viability based on trypan blue staining. More than 200 cells were counted per sample, with blue-stained cells considered non-viable and unstained cells considered alive.

### Measuring autophagy

Autophagy was assessed using the yeast autophagy marker GFP-ATG8, which is cleaved in the vacuole to release free GFP—a stable fragment detectable by fluorescence microscopy and Western blot analysis. Wild-type BY4741 and its isogenic *asc1Δ* strain were co-transformed with p*GAL-TDP-43* (*CEN*, *LEU2*) *or* p*GAL-FUS* (*CEN*, *HIS3*) and *pCUP1-GFP-ATG8* (*CEN, URA3*). Transformants grown on plasmid-selective glucose plates were replica-plated onto galactose-containing media supplemented with 1% raffinose and 50 µM CuSO₄, and incubated for 18–20 hours at 30 °C. Following trypan blue staining, autophagy was quantified as the fraction of live cells exhibiting fluorescence localized exclusively to the vacuole. This method was adapted from (Park *et al*. 2022). For Western blot analysis, cells were harvested and lysed after growth on plasmid-selective galactose media supplemented with 1% raffinose and 50 µM CuSO₄. Lysate preparation and immunoblotting were performed as previously described (Park *et al*. 2022). Autophagy was measured by calculating the ratio of cleaved GFP to uncleaved GFP-ATG8.

Protein expression levels of TDP-43 or FUS in BY4741 wild-type and *asc1Δ* strains were determined using α-TDP-43 (1:3000; Proteintech Group, Rosemont, IL, USA) or α-FUS antibody (1:3000; Proteintech Group, Rosemont, IL, USA), respectively and α-PGK1 antibody (1:5,000; OriGene Antibodies, Rockville, MD, USA) was used to detect Phosphoglycerate Kinase 1 as an internal loading control.

### Measuring TOROID formation

Toroid formation was measured as described previously (Park *et al*. 2024). Briefly, BY4741 (WT) and *asc1Δ* yeast strains carrying GFP-TOR1 were transformed with either p*GAL-TDP-43* (*CEN*, *URA3*) or empty vector. Cells were grown on selective galactose medium for 2 days and examined live with fluorescence microscopy using a FITC filter. TOROID foci were scored as punctate GFP-TOR1 structures, and the percentage of cells with foci was calculated relative to total live cells.

### Coimmunoprecipitation

To assess the physical interaction between TDP-43 and ASC1, BY4741 [*PIN*⁺] wild-type was transformed with either an empty vector (p2245, *pGAL-ccdB, CEN, LEU2*) or *pGAL-TDP-43* (p2368, *CEN, LEU2*). Transformants were grown in galactose-containing medium to induce TDP-43 expression, and cell lysates were prepared as previously described (Park *et al*. 2024).

Fresh transformants expressing TDP-43 were harvested after 18–20 hours of growth at 30 °C in galactose medium. Lysates were prepared and incubated for 10 minutes at 4 °C following the addition of Triton X-100 (final concentration 0.2%). Samples were centrifuged at 10,000×g for 10 minutes at 4 °C to obtain pre-cleared lysates.

For immunoprecipitation, 500 µL of pre-cleared lysate (2.0 mg/mL total protein) was incubated with 2 µL of α-TDP-43 antibody (Proteintech Group) for 2 hours on ice. Samples were then mixed with 50 µL of magnetic beads conjugated with immobilized G protein (Miltenyi Biotec Inc., Auburn, CA, USA) and incubated for 1 hour on ice. After washing to remove non-specifically bound proteins, co-immunoprecipitated proteins were eluted with hot sample buffer.

Eluted proteins were analyzed by SDS-PAGE and immunoblotting using α-TDP-43 (Proteintech Group, Rosemont, IL, USA) and α-ASC1 (MyBiosource Inc., San Diego, CA, USA) antibodies.

## Results

### Deletion of *ASC1* reduces TDP-43 but not FUS toxicity

Expression of TDP-43 in *S. cerevisiae* (BY4741 strain) significantly impairs cell growth. However, this toxicity is alleviated in the isogenic *asc*1Δ strain, despite a slight growth defect in *asc1Δ* cells in the absence of TDP-43 (Fig. 1A). Importantly, the frequency of dead cells caused by TDP-43 expression is significantly reduced in the *asc1Δ* background (Fig. 1B), indicating a protective effect. This relief of toxicity is not due to a change in the level of TDP-43 in the cell, as western blot analysis shows comparable expression in both wild-type and *asc*1Δ strains (Fig. 1C). In contrast, deletion of ASC1 does not mitigate FUS-induced toxicity, as growth inhibition and cell death remain unchanged in the presence of FUS (Fig. 1).

**Fig. 1.**
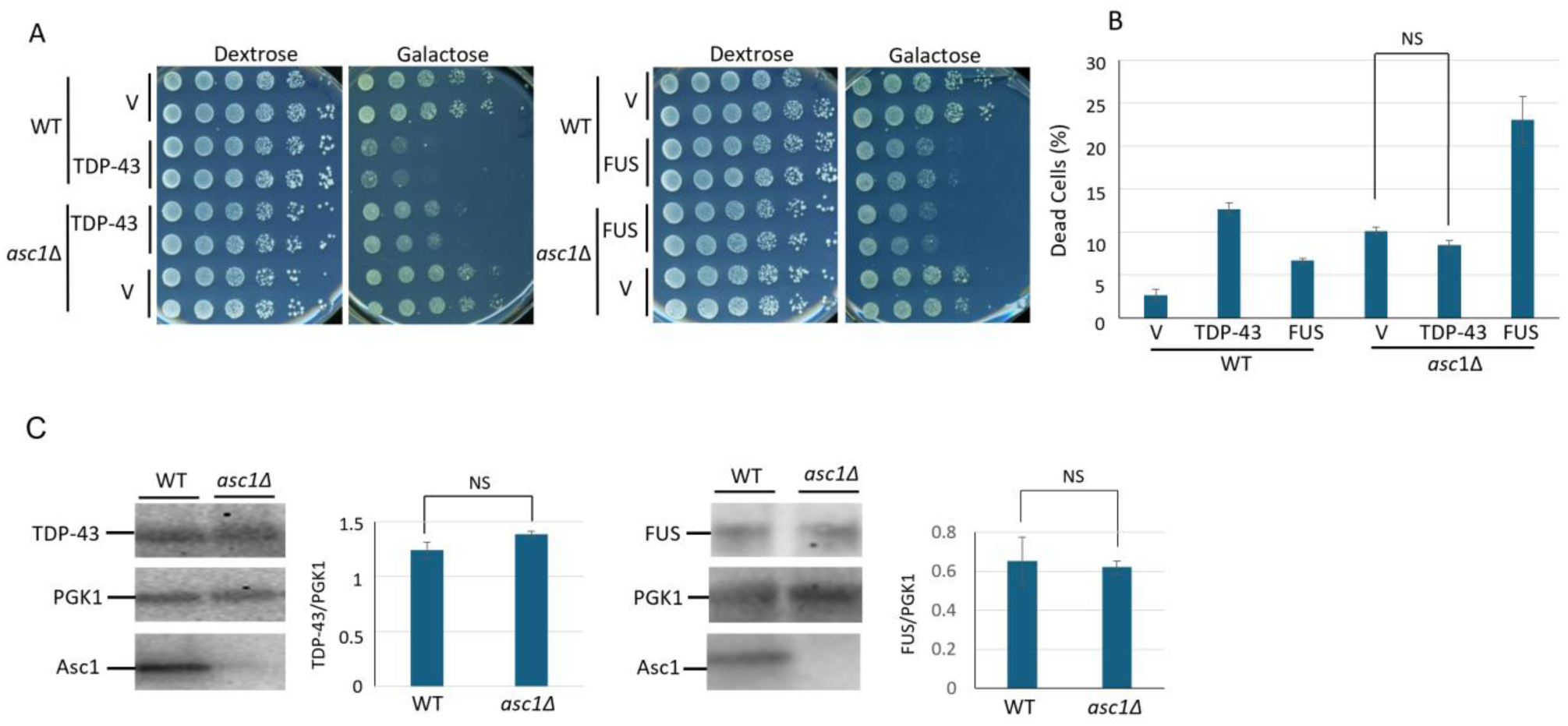
Deletion of *ASC1* partially rescues cells from TDP-43 toxicity but not FUS toxicity. A. Spot test showing growth of wild-type (WT) and *asc1Δ* strains transformed with empty vector (V), pGAL-TDP-43-YFP (TDP-43), or pGAL-FUS-YFP (FUS). Cultures grown overnight in plasmid-selective glucose media were concentrated to OD_600_=5, serially diluted 10-fold and spotted on non-inducing (Dextrose) and inducing (Galactose) synthetic plasmid-selective media. B. Strains described in A transformed with *pCUP-GFP-ATG8* (*URA3, CEN*) to monitor autophagy and grown overnight at 30°C in plasmid-selective galactose media with 50 µM Cu²⁺ to induce TDP-43 and GFP-ATG8. Cells were then stained with trypan blue to detect dead cells and the percentage dead cells among GFP-positive cells was quantified. Three sets of transformant for each were used and > 300 cells for each were counted. Error bars indicate standard error of the mean (SEM) and ** indicates p > 0.05. C. The levels of TDP-43 or FUS are unchanged in wild-type vs. *asc1Δ* strains. Western blot of the WT and *asc1Δ* strains described in A shows galactose induced TDP-43 or FUS and loading control, phosphoglycerate kinase (PGK1), levels. The bar graph shows data from three independent such Westerns. Error bars represent the SEM.

### Deletion of *ASC1* reduces the size and frequency of TDP-43 foci but they remain liquid-like and fail to stain with thioflavin T

Our previous work demonstrated that TDP-43 foci are dissolved by hexanediol and are not stained with thioflavin T, even in the presence of *pbp1Δ* and *tip41Δ,* which reduce TDP-43 toxicity (Park *et al*. 2024). Thus these foci are liquid-like rather than amyloid-like. However we found that *aah1Δ apt1Δ,* which also reduces TDP-43 toxicity shifted TDP-43 foci to be more amyloid like judged by altered responses to hexanediol, thioflavin T, and FRAP (Park *et al*. 2025). Deletion of *ASC1* falls in the category of *pbp1Δ* and *tip41Δ,* reducing TDP-43 toxicity without changing TDP-43 aggregate response to hexanediol or thioflavin T (Fig. 2A).

**Fig. 2.**
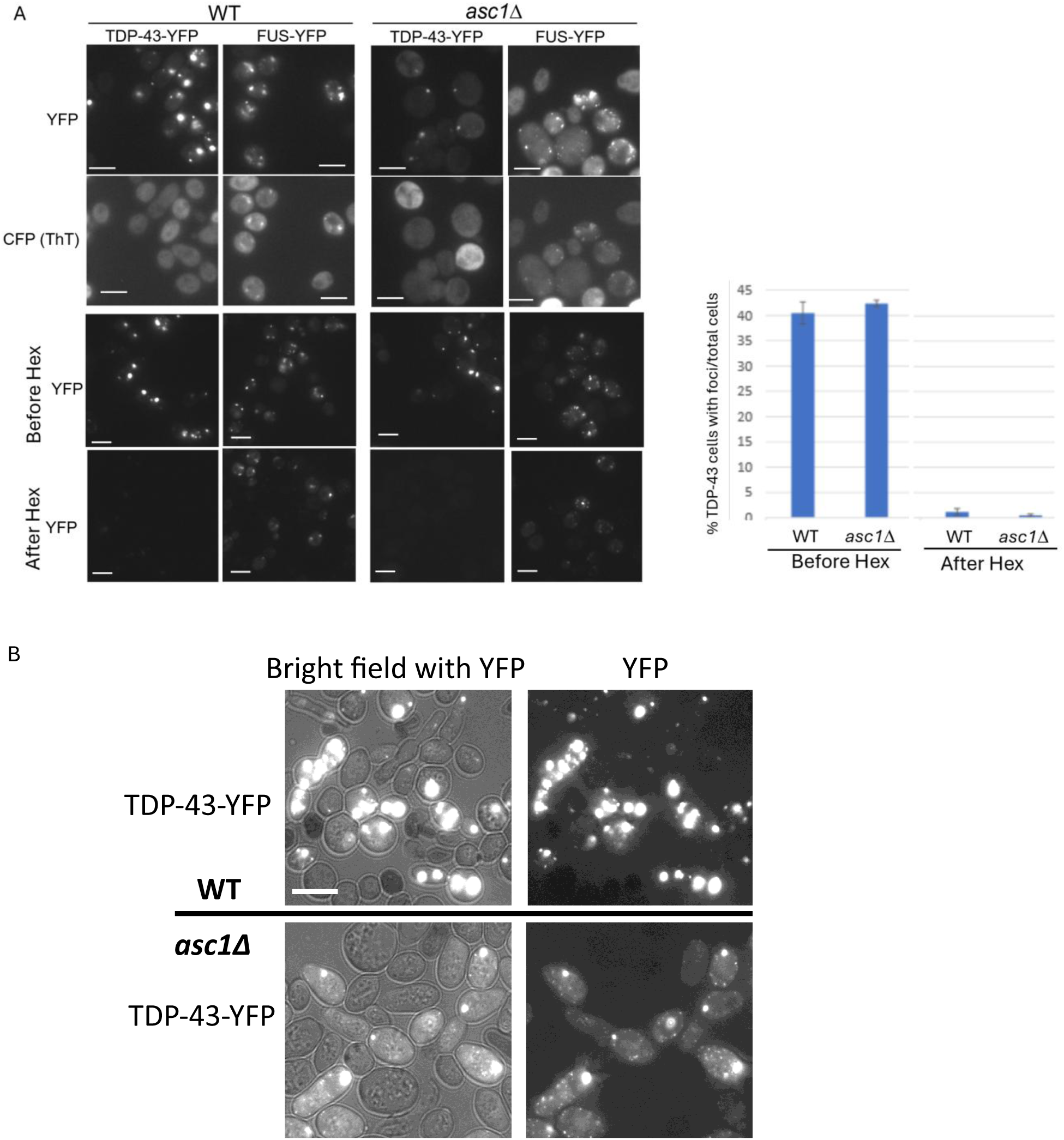
TDP-43 foci in *asc1Δ* strains remain liquid-like but have altered morphology. A. TDP-43-YFP foci don’t stain with thioflavin T and are dissolved by hexanediol in wild-type or *asc1Δ* strains. As reported previously, *asc1Δ* cells are larger than isogenic *ASC1* wild-type (WT) cells (Jorgensen *et al*. 2002). Scale bar is 10 µm. Upper panels: BY4741 (WT) and the isogenic *asc1Δ* strains transformed with *pGAL-TDP43-YFP (CEN, URA3)* or *pGAL-FUS-YFP* (*CEN, URA3*) were grown on galactose plasmid-selective plates. Cells expressing TDP-43-YFP or FUS-YFP were stained with Thioflavin T (ThT) and examined for YFP fluorescence or CFP fluorescence to assess the ThT staining. Lower panels and right: TDP-43-YFP foci were dissolved by hexanediol in both wild-type or *asc1Δ* strains. YFP fluorescence of cells described above expressing TDP-43-YFP was examined before (Before Hex) and after (After Hex) treatment with 10% 1,6-hexanadiol for 5 minutes that dissolves liquid-like aggregates. Three transformants each were tested. Bars show standard error of the mean. B. TDP-43 foci in *asc*1Δ strains are smaller and more numerous, with many tiny foci observed compared to WT strains.

### ASC1 is not sequestered into TDP-43 foci in yeast

Aggregates of TDP-43 in ALS patient spinal cord motor neurons were reported to co-aggregate with RACK1. This led to the hypothesis that TDP-43 aggregates sequester polysomes resulting in inhibition of protein synthesis leading to toxicity (Russo *et al*. 2017). Since TDP-43 is also toxic in yeast we asked if the RACK1 homolog, ASC1, was sequestered into TDP-43 foci in yeast, but found that it was not. Yeast cells expressing TDP-43-YFP from a *GAL* promoter were co-transformed with either *pPASC1-ASC1-mCFP* (*CEN*, native promoter) or *pGAL-ASC1-mCFP* (2*µ*, *GAL* promoter). In both cases, TDP-43 formed distinct cytoplasmic foci, while ASC1-mCFP remained diffusely distributed throughout the cytoplasm, showing no evidence of co-localization (Fig. 3A).

**Fig. 3.**
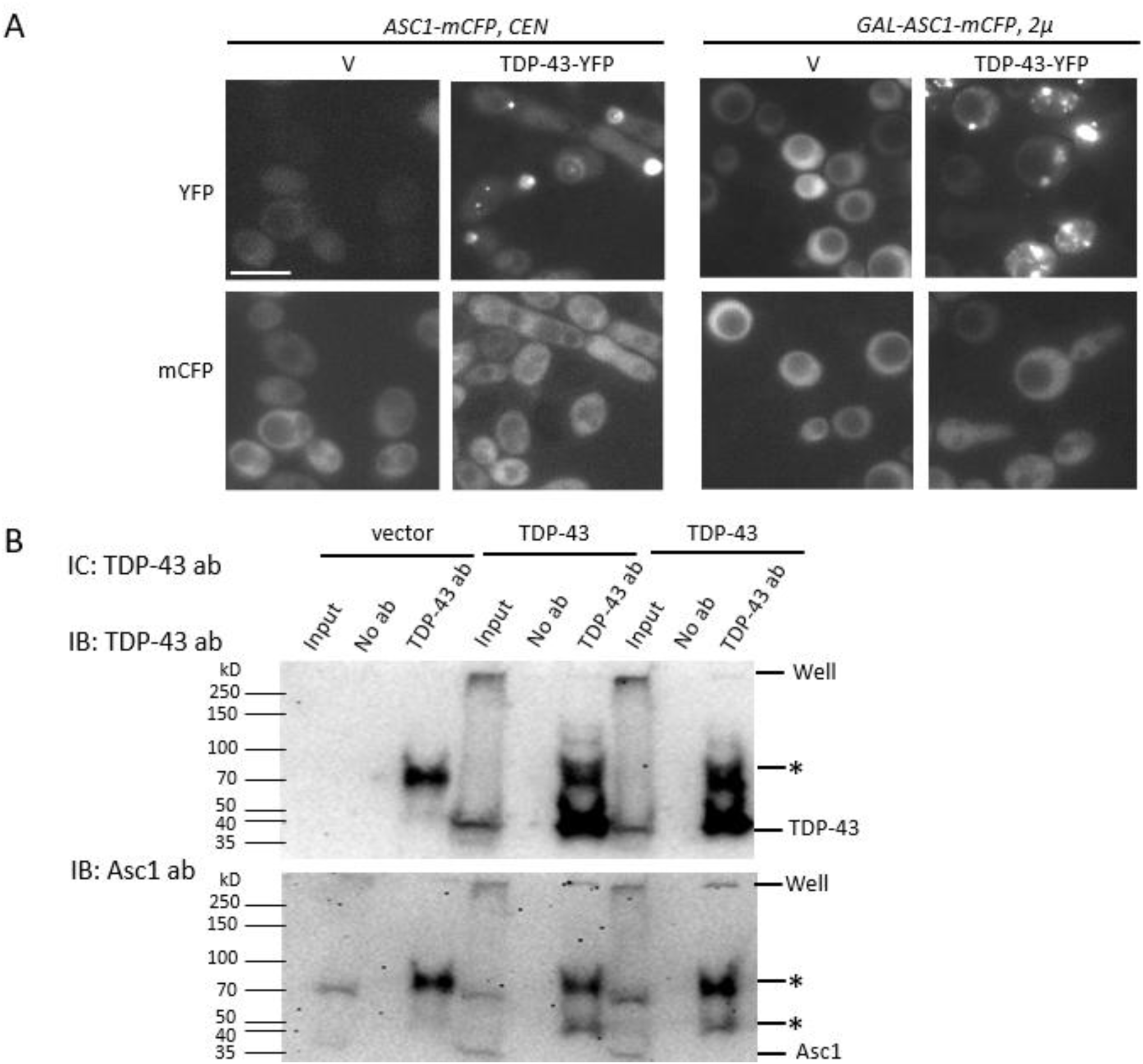
ASC1 is not sequestered into TDP-43 foci. A. ASC1 does not co-localize with TDP-43 foci in yeast. An *asc1Δ* strain was co-transformed with either YFP-tagged empty vector (V) or p*GAL-TDP-43-YFP, URA3, CEN* (TDP-43-YFP) and with either p*PASC1*-*ASC1-mCFP* (*ASC-mCFP*, CEN) or p*GAL-ASC1-mCFP* (*GAL-ASC1-mCFP*, *2µ*). Cells were patched on plasmid-selective galactose plates, grown overnight and, examined for fluorescence. The left panels show expression of YFP-tagged empty vector (V), and the right panels show YFP-tagged TDP-43 (TDP-43-YFP). Cerulean-tagged ASC1 does not form foci. Scale bar, 10 μm. B. ASC1 does not co-immunoprecipitate with TDP-43. TDP-43 was immunocaptured (IC) with TDP-43 antibody (TDP-43 ab) from cells transformed either an empty vector (vector, p2245, *pGAL-ccdB, CEN, LEU2*) or TDP-43 (p2368, *pGAL-TDP-43, CEN, LEU2*). Western blots of input, no-antibody control, and TDP-43 immunoprecipitates were probed sequentially with TDP-43 and ASC1 antibodies. Non-specific bands are marked with *.

To further assess physical interaction, we also asked if ASC1 would co-immunoprecitate with TDP-43. TDP-43 was immunocaptured from cells transformed with either an empty vector control (vector) or pGAL-TDP-43 (TDP-43). Western blots of input, no antibody control, and TDP-43 immunoprecipitates revealed ASC1 only in the input samples, with no detectable ASC1 in the TDP-43 pulldown (Fig. 3B). Thus while TDP-43 and RACK1 interact in a two-hybrid assays (Adams *et al*. 2011; Dengjel *et al*. 2012) (http://www.interactome-atlas.org/), ASC1 was neither recruited into TDP-43 foci nor co-immunoprecipitated with TDP-43 in yeast. Thus, there is no evidence in yeast for the hypothesis that ASC1 and associated polysomes are captured by TDP-43 aggregates resulting in inhibition of translation and causing TDP-43 toxicity.

### Deletion of *ASC1* prevents TDP-43 from inhibiting autophagy

One proposed mechanism for TDP-43 toxicity in yeast is its inhibition of autophagy (Park *et al*. 2022). Furthermore, previously identified TDP-43 toxicity modifiers, *pbp1Δ*, *tip41Δ* and *aah1Δ apt1Δ* each alleviate TDP-43-induced autophagy inhibition (Park *et al*. 2024). Interestingly, deletion of *ASC1* has been shown to enhance autophagy (Dengjel *et al*. 2012). We now find that in *asc1Δ* strains, overexpression of TDP-43 no longer inhibits autophagy below the level observed in wild-type cells without TDP-43 (Fig. 4). This result supports the hypothesis that TDP-43 toxicity results from its inhibition of autophagy. In contrast, FUS does not inhibit autophagy in wild-type cells nor, in *asc1Δ,* cells does FUS reduce autophagy below that seen in wild-type cells.

**Fig. 4.**
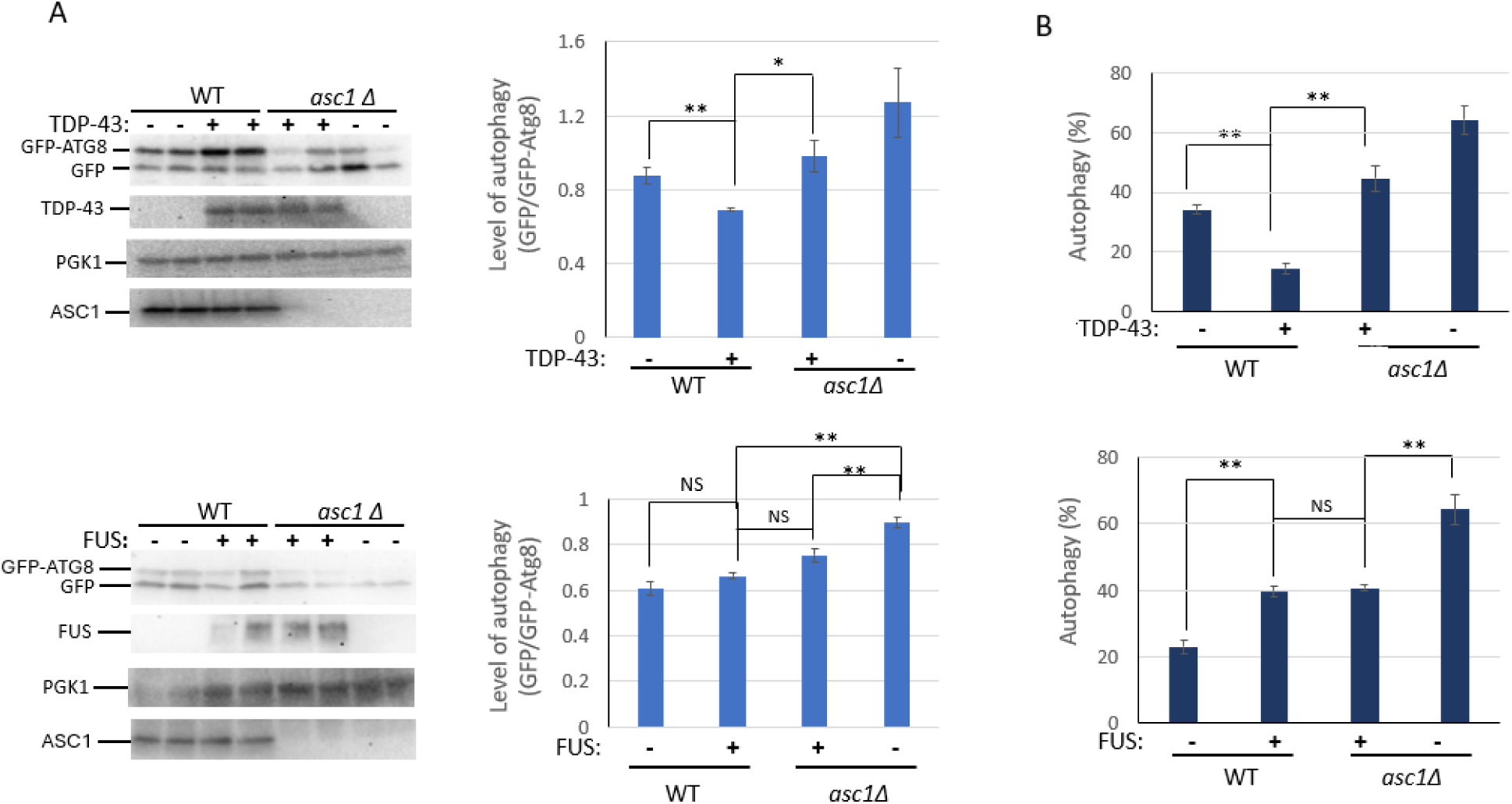
The *asc1Δ* deletion enhances autophagy and prevents TDP-43 from suppressing autophagy below wild-type base line levels. A. Autophagy measured by Western blot analysis of the breakdown of GFP-ATG8. BY4741 (WT) and the isogenic *asc1Δ* deletion were co-transformed with *pGAL-TDP43 (CEN, LEU2)* or *pGAL-FUS (CEN, HIS3)* along with p*CUP1-GFP-ATG8 (CEN, URA3)*. Cells were grown on 2% galactose plasmid-selective plates supplemented with 1% raffinose, the required amino acids, and 50 μM copper sulfate for 16 hours at 30oC before being harvested for Western blotting. One representative blot is shown (left), and quantification from three independent experiments is presented in the bar graph (right). **B.** Autophagy measured microscopically by GFP-ATG8 localization. The same transformants described in (A) were examined for autophagy by determining the subcellular localization of GFP-ATG8. Autophagy was quantified as the fraction of cells showing GFP fluorescence exclusively in the vacuole among all live cells with detectable GFP signal. Autophagy was increased in the *asc1Δ* cells whether or not TDP-43 was expressed above that seen in wild-type cells without TDP-43. Error bars represent the SEMs from three independent transformants, with 250-550 cells analyzed per transformant. ** indicates p < 0.01 (paired two-tailed t-test).

(Fig. 4).

### Deletion of *ASC1* lowers the frequency of TOROIDs

We previously demonstrated that TDP-43 toxicity modifiers, *pbp1Δ*, *tip41Δ* and *aah1Δ apt1Δ,* each reduce the inhibition of TOROID formation caused by TDP-43 overexpression (Park *et al*. 2024; Park *et al*. 2025). In contrast, while *asc1Δ* decreases TDP-43 toxicity and inhibition of autophagy, it does not increase the frequency of TOROIDs in the presence of TDP-43. Rather, *asc1Δ* decreases TOROID formation whether or not TDP-43 is expressed (Fig. 5). Since *asc1Δ* enhances autophagy (Fig. 4), this reduction in TOROID formation was unexpected and suggests that ASC1 influences TOROID dynamics through a distinct mechanism.

**Fig. 5.**
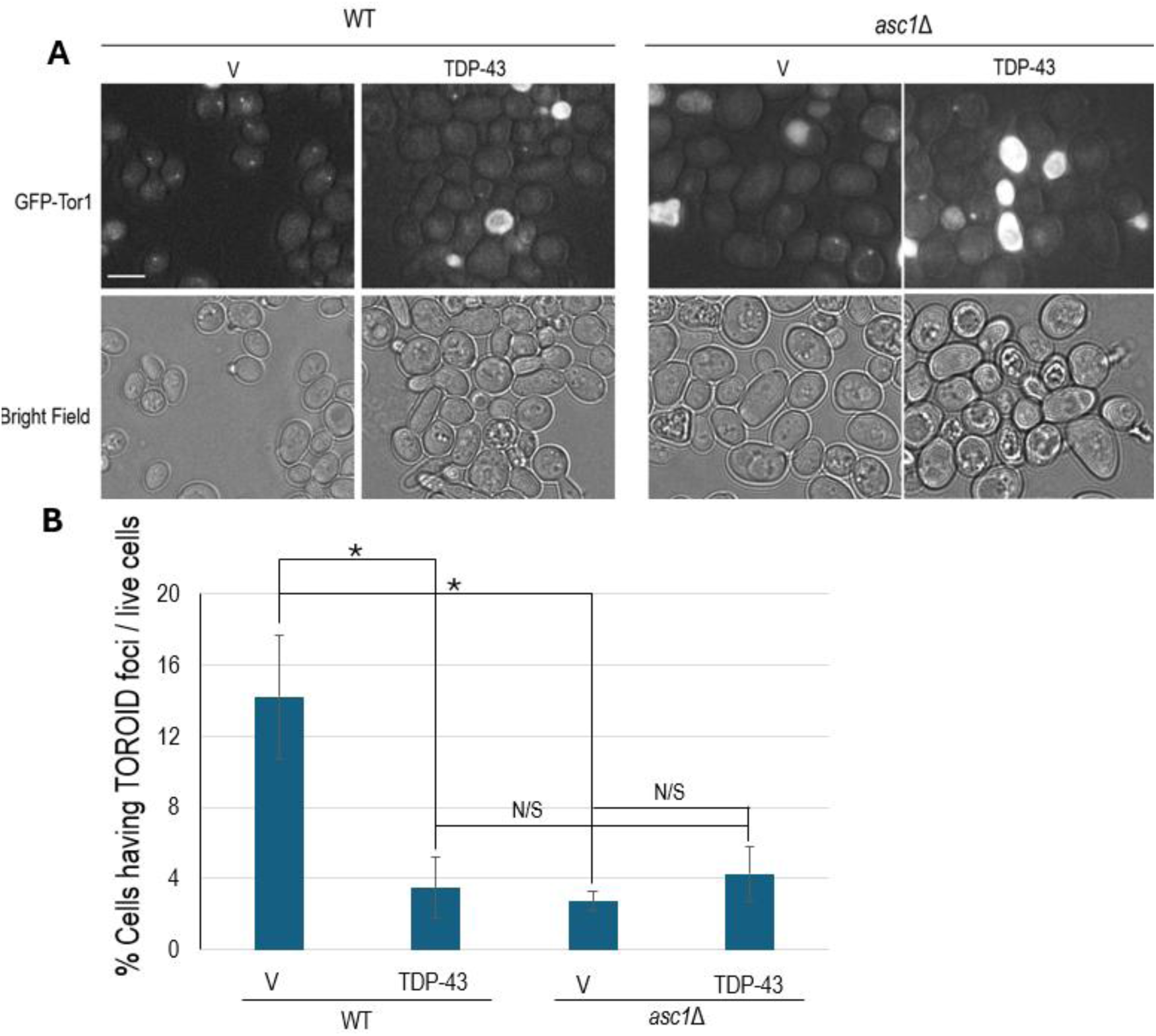
Deletion of *ASC1* inhibits the frequency of TOROIDs whether or not TDP-43 is expressed. A. Shown are fluorescent images of endogenously tagged *TOR1* (*GFP-TOR1*) [*PIN*^+^] BY4741 cells (WT) or in *asc1Δ* cells. Cells were transformed with untagged TDP-43 expressed under the *GAL* promoter in *pGAL-TDP43* (TDP-43, *CEN, URA3*) or empty vector *pGAL-ccdB (*V, *CEN, URA3*)), grown on selective galactose media for 2 days, examined with a FITC filter and photographed (upper). Size bar is 10 µm. B. Error bars represent SEMs calculated from three independent transformants. Statistical analysis using paired two-tailed t-tests yielded the following p-values: WT vector vs. TDP-43, 0.0494; BY(WT) vector vs. Δasu9 vector, 0.0305; Δasu9 vector vs. TDP-43, 0.4012. Asterisks (*) indicate statistically significant differences (p < 0.05); “NS” denotes no significant difference.

## Discussion

*Saccharomyces cerevisiae* continues to serve as a valuable model for studying mechanisms underlying neurodegenerative disease. Deletion of *ASC1*, the yeast ortholog of *RACK1* (Gerbasi et al. 2004), reduced TDP-43 toxicity in yeast just as reducing *RACK1* expression in *Drosophila melanogaster* decreases TDP-43 toxicity (Zhao *et al*. 2023). Additionally, deletion of *ASC1/RACK1* alters the morphology of TDP-43 foci. In mammalian cells cytoplasmic aggregation of TDP-43 lacking its nuclear localization signal is reduced by knockdown of RACK1 and TDP-43 actually returns to the nucleus (Zhao *et al*. 2023). In the yeast model, *asc1Δ* reduces the size of TDP-43 foci, although TDP-43 remains in the cytoplasm.

In mammalian cell culture, both TDP-43 and FUS were shown to inhibit global protein synthesis, which was partially prevented by knockdown of *RACK1*. Additionally, TDP-43 and FUS co-aggregated with RACK1, a 40S ribosomal protein. This led to the hypothesis that toxicity results from sequestration of polysomes by TDP-43 and FUS and the resulting translational inhibition (Russo *et al*. 2017; Zhao *et al*. 2023).

We tested this hypothesis in yeast and found that while *asc1Δ* reduced TDP-43 toxicity, it did not affect FUS toxicity. Furthermore, ASC1 remained diffuse in yeast cytoplasm expressing TDP-43, despite the presence of TDP-43 foci. Also, ASC1 did not co-immunoprecipitate with TDP-43.

These observations are inconsistent with a model in which TDP-43 and FUS sequester polysomes via ASC1/RACK1 and inhibit translation as the primary mechanism of toxicity. We previously showed that TDP-43 expression inhibits autophagy (Park *et al*. 2022) and TOROID formation (Park *et al*. 2024) in yeast. Now we show that *asc1Δ* rescues cells from the reduction in autophagy without rescuing TOROID formation. Interestingly, while TDP-43 inhibits autophagy, FUS expression doesn’t (Fig. 4). Thus we propose that effects on autophagy underlie TDP-43 but not FUS toxicity, and that inhibition of autophagy by TDP-43 may secondarily reduce translation.

We confirm that deletion of *ASC1* enhances autophagy even in the absence of TDP-43 (Dengjel *et al*. 2012), indicating that ASC1 is a negative regulator of autophagy. If reduced autophagy contributes to TDP-43 toxicity, then its upregulation in *asc1Δ* cells explains the observed rescue. Furthermore, increased autophagy caused by reduction in RACK1/ASC1 could explain the observed absence of large TDP-43 foci (Huang *et al*. 2020).

Canonical autophagy is enhanced in yeast when TOROIDS form and sequester TORC1, thereby relieving TORC1 inhibition of autophagy (Noda 2017; Prouteau *et al*. 2017). Interestingly, although *asc1Δ* enhanced autophagy, it also reduced TOROID formation (Fig. 5), suggesting activation of a non-canonical autophagy pathway that works independently of TORC1. In support of this, knockdown of RACK1 in mammalian cells was shown to induce non-canonical autophagy by selective regulation of translation initiation (Kim *et al*. 2017; Silvestri *et al*. 2025).

We hypothesize that *asc1Δ* induces non-canonical autophagy, and that cells attempt to compensate for this by dissolving TOROIDS to reactivate TORC1 and suppress canonical autophagy. However, because TORC1 does not inhibit non-canonical autophagy, autophagy remains elevated despite the reduced TOROID number.

In contrast, to *asc1Δ* other yeast modifiers of TDP-43 toxicity do not alter autophagy in the absence of TDP-43. Also their effects on TOROID formation align with changes in autophagic activity pointing to a canonical autophagy pathway (Park *et al*. 2022; Park *et al*. 2024; Park *et al*. 2025).

Our results highlight autophagy, rather than polysome sequestration, as a central mechanism in the toxicity of TDP-43 and rescue by reduced ASC1/RACK1.

## Acknowledgments

We thank Irina Derkatch for helpful suggestions and Dr. Robbie Joséph Loewith for generously providing the pSK108 construct (Prouteau et al. 2017) and Dr. Aaron Gitler for sharing plasmids on Addgene.

## Data and reagent availability

The authors affirm that all data necessary for confirming the conclusions of the article are present within the article, figures, and tables. Strains and plasmids are available upon request.

## Literature cited

Adams, D. R., D. Ron and P. A. Kiely, 2011 RACK1, A multifaceted scaffolding protein: Structure and function. Cell Commun Signal 9: 22.

Armakola, M., M. P. Hart and A. D. Gitler, 2011 TDP-43 toxicity in yeast. Methods 53: 238–245.

Becker, L. A., B. Huang, G. Bieri, R. Ma, D. A. Knowles et al., 2017 Therapeutic reduction of ataxin-2 extends lifespan and reduces pathology in TDP-43 mice. Nature 544: 367–371.

Capitini, C., G. Fani, M. Vivoli Vega, A. Penco, C. Canale et al., 2021 Full-length TDP-43 and its C-terminal domain form filaments in vitro having non-amyloid properties. Amyloid 28: 56–65.

Cascella, R., M. Banchelli, S. Abolghasem Ghadami, D. Ami, M. C. Gagliani et al., 2023 An in situ and in vitro investigation of cytoplasmic TDP-43 inclusions reveals the absence of a clear amyloid signature. Ann Med 55: 72–88.

Dengjel, J., M. Hoyer-Hansen, M. O. Nielsen, T. Eisenberg, L. M. Harder et al., 2012 Identification of autophagosome-associated proteins and regulators by quantitative proteomic analysis and genetic screens. Mol Cell Proteomics 11: M111 014035.

Dormann, D., and C. Haass, 2011 TDP-43 and FUS: a nuclear affair. Trends Neurosci 34: 339–348.

Fushimi, K., C. Long, N. Jayaram, X. Chen, L. Li et al., 2011 Expression of human FUS/TLS in yeast leads to protein aggregation and cytotoxicity, recapitulating key features of FUS proteinopathy. Protein Cell 2: 141–149.

Gerbasi, V. R., C. M. Weaver, S. Hill, D. B. Friedman and A. J. Link, 2004 Yeast Asc1p and mammalian RACK1 are functionally orthologous core 40S ribosomal proteins that repress gene expression. Mol Cell Biol 24: 8276–8287.

Gispert, S., A. Kurz, S. Waibel, P. Bauer, I. Liepelt et al., 2012 The modulation of Amyotrophic Lateral Sclerosis risk by ataxin-2 intermediate polyglutamine expansions is a specific effect. Neurobiol Dis 45: 356–361.

Gitler, A. D. a. E. A. C., 2012 Modulators of TDP-43 Mediated Toxicity and Methods of Use Thereof for Identifying Agents Having Efficacy for the Treatment and Prevention of Proteinopathies., pp., USA.

Gross, A. S., A. Zimmermann, T. Pendl, S. Schroeder, H. Schoenlechner et al., 2019 Acetyl-CoA carboxylase 1-dependent lipogenesis promotes autophagy downstream of AMPK. J Biol Chem 294: 12020–12039.

Huang, C., S. Yan and Z. Zhang, 2020 Maintaining the balance of TDP-43, mitochondria, and autophagy: a promising therapeutic strategy for neurodegenerative diseases. Transl Neurodegener 9: 40.

Johnson, B. S., J. M. McCaffery, S. Lindquist and A. D. Gitler, 2008 A yeast TDP-43 proteinopathy model: Exploring the molecular determinants of TDP-43 aggregation and cellular toxicity. Proc Natl Acad Sci U S A 105: 6439–6444.

Jorgensen, P., J. L. Nishikawa, B. J. Breitkreutz and M. Tyers, 2002 Systematic identification of pathways that couple cell growth and division in yeast. Science 297: 395–400.

Ju, S., D. F. Tardiff, H. Han, K. Divya, Q. Zhong et al., 2011 A yeast model of FUS/TLS-dependent cytotoxicity. PLoS Biol 9: e1001052.

Kim, H. D., E. Kong, Y. Kim, J. S. Chang and J. Kim, 2017 RACK1 depletion in the ribosome induces selective translation for non-canonical autophagy. Cell Death Dis 8: e2800.

Ma, X. R., M. Prudencio, Y. Koike, S. C. Vatsavayai, G. Kim et al., 2022 TDP-43 represses cryptic exon inclusion in the FTD-ALS gene UNC13A. Nature 603: 124–130.

Noda, T., 2017 Regulation of Autophagy through TORC1 and mTORC1. Biomolecules 7.

Park, S., S. K. Park, P. Blair and S. W. Liebman, 2025 An adenine model of inborn metabolism errors alters TDP-43 aggregation and reduces its toxicity in yeast revealing insights into protein misfolding diseases. Microb Cell 12: 119–130.

Park, S., S. K. Park and S. W. Liebman, 2024 Expression of Wild-Type and Mutant Human TDP-43 in Yeast Inhibits TOROID (TORC1 Organized in Inhibited Domain) Formation and Autophagy Proportionally to the Levels of TDP-43 Toxicity. Int J Mol Sci 25.

Park, S. K., S. Park and S. W. Liebman, 2022 TDP-43 Toxicity in Yeast Is Associated with a Reduction in Autophagy, and Deletions of TIP41 and PBP1 Counteract These Effects. Viruses 14.

Prouteau, M., A. Desfosses, C. Sieben, C. Bourgoint, N. Lydia Mozaffari et al., 2017 TORC1 organized in inhibited domains (TOROIDs) regulate TORC1 activity. Nature 550: 265–269.

Russo, A., R. Scardigli, F. La Regina, M. E. Murray, N. Romano et al., 2017 Increased cytoplasmic TDP-43 reduces global protein synthesis by interacting with RACK1 on polyribosomes. Hum Mol Genet 26: 1407–1418.

Shenoy, J., A. Lends, M. Berbon, M. Bilal, N. El Mammeri et al., 2023 Structural polymorphism of the low-complexity C-terminal domain of TDP-43 amyloid aggregates revealed by solid-state NMR. Front Mol Biosci 10: 1148302.

Silvestri, F., R. Montuoro, E. Catalani, F. Tilesi, D. Willems et al., 2025 eIF3d specialized translation requires a RACK1-driven eIF3d binding to 43S PIC in proliferating SH-SY5Y neuroblastoma cells. Cell Signal 125: 111494.

Wolf, A. S., and E. J. Grayhack, 2015 Asc1, homolog of human RACK1, prevents frameshifting in yeast by ribosomes stalled at CGA codon repeats. RNA 21: 935–945.

Zeng, Y., A. Lovchykova, T. Akiyama, C. Liu, C. Guo et al., 2024 TDP-43 nuclear loss in FTD/ALS causes widespread alternative polyadenylation changes. bioRxiv.

Zhao, B., C. M. Cowan, J. A. Coutts, D. D. Christy, A. Saraph et al., 2023 Targeting RACK1 to alleviate TDP-43 and FUS proteinopathy-mediated suppression of protein translation and neurodegeneration. Acta Neuropathol Commun 11: 200.

